# Reference-free Cell-type Annotation for Single-cell Transcriptomics using Deep Learning with a Weighted Graph Neural Network

**DOI:** 10.1101/2020.05.13.094953

**Authors:** Xin Shao, Haihong Yang, Xiang Zhuang, Jie Liao, Yueren Yang, Penghui Yang, Junyun Cheng, Xiaoyan Lu, Huajun Chen, Xiaohui Fan

**Author notes:** These authors have contributed equally to this work (Xin Shao, Haihong Yang).

## Abstract

Advances in single-cell RNA sequencing (scRNA-seq) have furthered the simultaneous classification of thousands of cells in a single assay based on transcriptome profiling. In most analysis protocols, single-cell type annotation relies on marker genes or RNA-seq profiles, resulting in poor extrapolation. Here, we introduce scDeepSort (https://github.com/ZJUFanLab/scDeepSort), a reference-free cell-type annotation tool for single-cell transcriptomics that uses a deep learning model with a weighted graph neural network. Using human and mouse scRNA-seq data resources, we demonstrate the feasibility of scDeepSort and its high accuracy in labeling 764,741 cells involving 56 human and 32 mouse tissues. Significantly, scDeepSort outperformed reference-dependent methods in annotating 76 external testing scRNA-seq datasets, including 126,384 cells (85.79%) from ten human tissues and 134,604 cells from 12 mouse tissues (81.30%). scDeepSort accurately revealed cell identities without prior reference knowledge, thus potentially providing new insights into mechanisms underlying biological processes, disease pathogenesis, and disease progression at a single-cell resolution.

## Introduction

Recent advancements in single-cell RNA sequencing (scRNA-seq) that permit the identification of various cell types based on transcriptomics at single-cell resolution have facilitated our understanding of the heterogeneity of cellular phenotypes and their composition within complex tissues^1,2^. In the data processing protocols of scRNA-seq experiments, cell-type annotation is a vital step for subsequent analysis^3^. Cell type identification is commonly performed by mapping differentially expressed genes at the level of pre-computed clusters with prior knowledge of cell markers like scCATCH^4^. Another cell-based annotation strategy tries to compare the similarities between single cell and reference database of bulk or single-cell RNA-seq profiles to determine potential cellular identities. Several methods including SingleR^5^, CHETAH^6^, scMap^7^ and scHCL^8^ belong to this category. An increasing number of machine learning-based annotation approaches—including CellAssign^9^ and Garnett^10^—have emerged as well in recent years. Such methods rely heavily on references, namely RNA-seq profiles with known cell types or known cell marker genes as prior knowledge, severely limiting the extrapolation of these methods. Still, accurate cell-type annotation for single-cell transcriptomic data remains a great challenge.

Fortunately, recent advances in deep learning have enabled major progress in the ability of artificial intelligence techniques to integrate big data, incorporate existing knowledge, and learn arbitrarily complex relationships^11,12^. Given the state-of-the-art accuracy deep learning has achieved in numerous prediction tasks, it has been increasingly used in biological research^13^ and biomedical applications^14^ such as drug discovery^15^, medical diagnosis^16^, as well as various-omics tasks^17^. For example, Kermany et al. combined optical coherence tomography images with a deep-learning framework and developed an efficient diagnostic tool to support the clinical decision for patients with blinding retinal diseases^18^. Chaudhary et al. established a deep learning-based model using RNA-seq, miRNA-seq and methylation data of 360 hepatocellular carcinoma (HCC) patients to help predict patient survival^19^. Graph neural networks (GNNs) are connectionist models which capture the graph dependence through message passing between the graph nodes. Unlike standard neural networks, GNNs retain a state that represents information from its neighborhood with arbitrary depth, which have demonstrated ground-breaking performance on many learning tasks^20^. Moreover, recent published large-scale scRNA-seq resources have provided the foundation for a GNN-based deep learning model that can execute challenging prediction tasks.

In this study, we designed a reference-free cell-type annotation method called scDeepSort, based on a weighted GNN framework, which addresses this challenge (see Fig. 1 for an overview). Briefly, scDeepSort is learned by a weighted GNN model for supervised training on recently published the most comprehensive single-cell transcriptomics atlases—a human cell landscape (HCL^8^) consisting of 56 tissues and 562,977 cells and a mouse cell atlas (MCA^21^) consisting of 32 tissues and 201,764 cells (Supplementary Table S1). scDeepSort showed high accuracy in cell type identification, significantly outperforming four current excellent methods—CellAssign, Garnett, SingleR, and scMap—on 76 external human and mouse testing datasets including 126,384 cells from ten human tissues and 134,604 cells from 12 mouse tissues. The present results indicated that scDeepSort can help scientists rapidly annotate a single cell with an accurate cell label without prior reference knowledge, i.e., markers or RNA-seq profiles, which may tremendously facilitate scRNA-seq studies and provide novel insights into the mechanisms underlying biological processes and disease pathogenesis and progression.

**Fig. 1.**
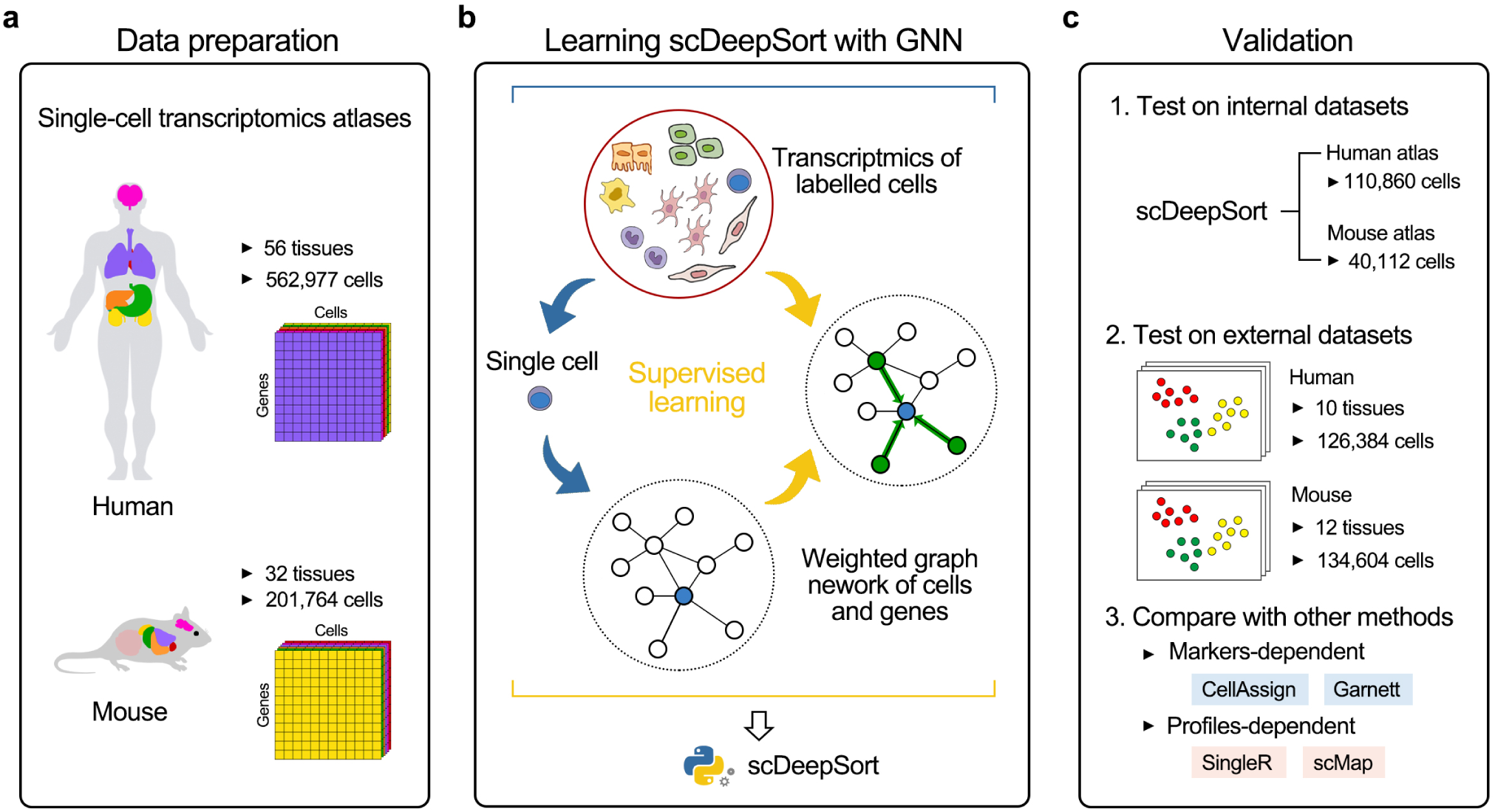
General conceptual framework and validation of scDeepSort. **a**) Human and mouse single-cell transcriptomics atlases were curated from HCL and MCA as the underlying data for training scDeepSort. The human and mouse atlases include 562,977 cells from 56 tissues and 201,764 cells from 32 tissues, respectively. **b**) For each cell, a graph network was constructed of this cell, its genes and neighboring cells for supervised learning scDeepSort with known cell labels from the transcriptomic atlases. **c**) Internal human and mouse atlases datasets and external testing datasets of single-cell transcriptomics data involving multiple tissues were employed to test the performance of scDeepSort. Markers- and profiles-dependent annotation methods (CellAssign, Garnett, SingleR and scMap) were compared with scDeepSort on human and mouse external testing datasets.

## Results

### General description of scDeepSort

In brief, we applied a supervised deep learning model based on a weighted GNN framework to build the scDeepSort model with underlying data of human and mouse single-cell transcriptomics atlases. First, dense representations for cells and genes were obtained with dimensionality reduction methods initialize fixed-size node embeddings, since the single-cell transcriptomics data are usually sparse matrices. Principal component analysis (PCA) was used to extract dense representations for gene nodes from the cell-gene data matrix, and cell node representations were then calculated by the weighted sum of gene node representations and the cell–gene data matrix. Then, an undirected and weighted graph containing cell nodes and gene nodes was constructed from an adjacent weighted matrix by taking the gene expression as the weighted edges between cells and genes to model the intrinsic geometric information, which constitutes scDeepSort’s first embedding layer (Fig. 2).

**Fig. 2.**
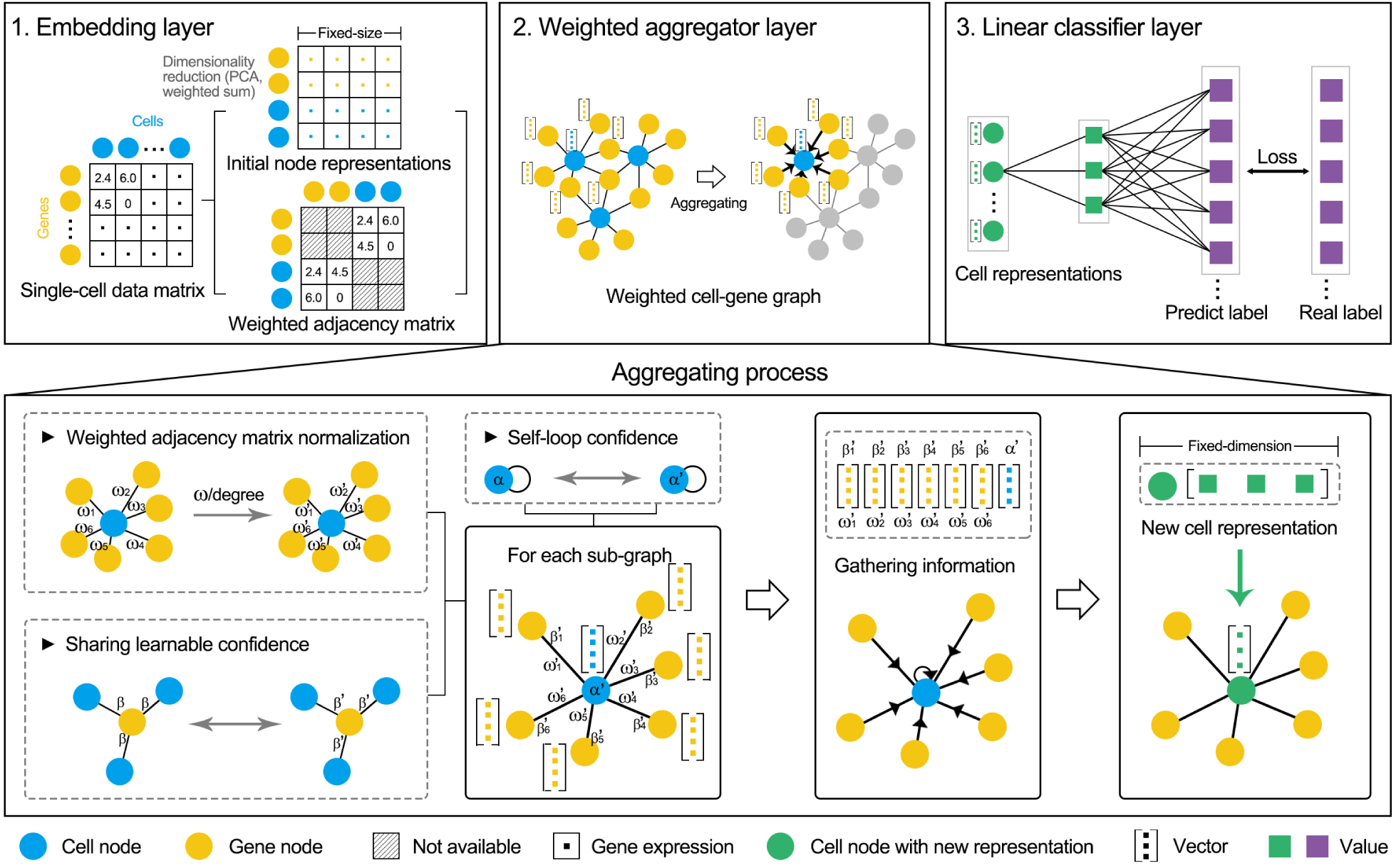
The weighted GNN algorithm of scDeepSort. The algorithm consists of embedding, weighted graph aggregator and linear classifier layers. The embedding layer stores the graph node representations and is freezed during training, wherein dimensionality reduction methods (PCA and weighted sums) were used to generate the initial fix-size node representations and the gene expression for each cell was regarded as the weighted edge between cells and genes forming a weighted adjacent matrix. In the weighted aggregator layer, a self-loop confidence for each cell node and a learnable sharing confidence for each gene node were incorporated into the weighted cell-gene graph. For each cell-centered subgraph, weighted edges were normalized and node itself and neighborhood were then gathered to generate a new cell node representation during aggregation. The linear classifier layer categorizes the final cell state representation as a predefined cell type.

**Fig. 3.**
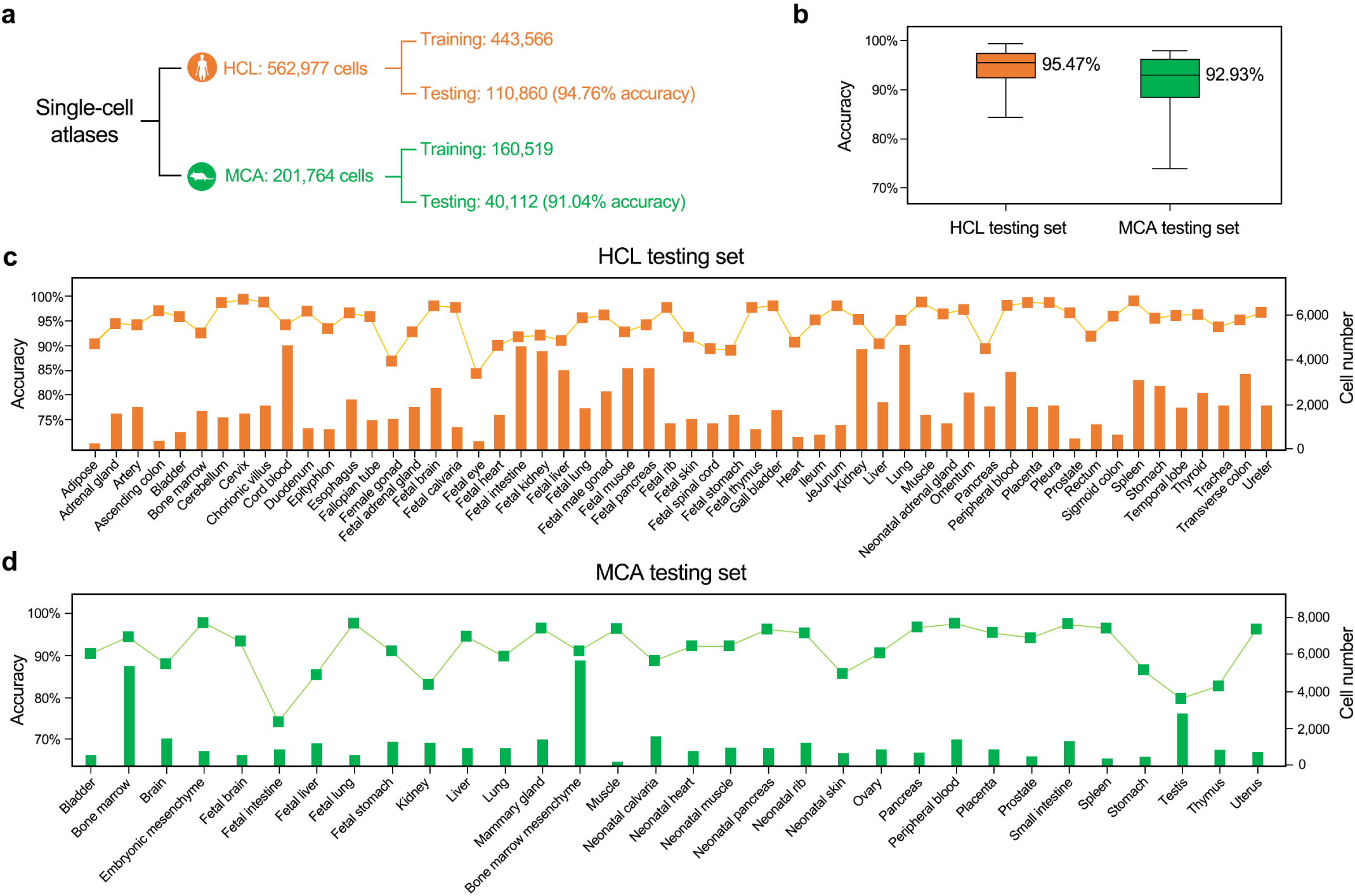
scDeepSort performance on internal datasets. **a**) Statistics of the HCL and MCA training and testing sets, as well as the accuracy on all testing cells. **b**) Boxplots show the distribution of accuracies on each tissue for HCL and MCA testing sets. The median accuracy is labelled beside the corresponding box. **c**) Accuracy of scDeepSort in annotating the HCL testing set for 56 tissues (line chart). The bar chart shows the number of test cells for each tissue. **d**) Accuracy of scDeepSort in annotating the MCA testing set for 32 tissues.

In detail, the scDeepSort model’s architecture consists of three components: the embedding layer, the weighted graph aggregator layer and the linear classifier layer (Fig. 2). The embedding layer stores representations of graph nodes, which are initialized as previously described and are freezed during training. The weighted graph aggregator layer uses inductive learning to ascertain graph structure information; in this layer, GraphSAGE^22^ was applied as the backbone GNN framework and heavily modified in some aspects. First, a normalized weighted adjacency matrix that is quite different from the adjacent matrix used in conventional GNN models was proposed and applied to the weighted cell-gene graph for each cell node, in consideration of the wide difference of expression level and pattern for different genes and cells. Second, a learnable sharing confidence for each gene node was incorporated into the weighted graph in order to avoid batch effect and solve the missing value issues. Third, a self-loop confidence was added to the weighted graph for each cell node. The weighted graph aggregator layer generates a linear separable feature space for cells. The final linear classifier layer classifies the final cell state representation produced from the weighted graph aggregator layer into one of the predefined cell type categories.

During training for each cell-centered subgraph, the cell node representation, along with its connected gene node representations from the embedding layer, was extracted and then aggregated on the weighted graph aggregator layer by gathering information about the neighborhood and itself, which produced a new cell representation for each cell. Once a label is predicted by the linear classifier layer, the loss between this prediction and the correct label is computed and then used to update the parameters of three layers until convergence (Fig. 2).

### Performance on internal datasets

In this study, a total of 562,977 human cells from 56 tissues and 201,746 mouse cells from 32 tissues which were collected from recently published HCL and MCA were curated to construct single-cell transcriptomic atlases (Supplementary Table S1). For each cell type, cells numbering fewer than 5‰ of the total cells in each tissue were dropped. For each tissue in the human and mouse atlases, cells of various types were first merged and randomly divided into training and testing sets, ensuring that the ratio of training and testing cells was set to 8:2 for each cell type. In total, this process generated 443,566 training cells and 110,860 testing cells from the HCL, 160,519 training cells and 40,112 testing cells from the MCA (Fig. 2a). After supervised learning on training sets, we evaluated the scDeepSort’s performance on HCL and MCA testing sets.

scDeepSort assigned labels with a 94.76% accuracy on 110,860 human testing cells and a 91.04% accuracy on 40,112 mouse testing cells (Fig. 2a). Although the accuracies were only up to 80% on mouse fetal brain cells and testis cells, the median accuracies across 56 human tissues and 32 mouse tissues reached 95.47% and 92.23% (Fig. 2b). Across 56 human tissues and 32 mouse tissues, scDeepSort annotated the most cells, with the accuracy ranging from 84.28% to 99.25% and from 73.88% to 97.88%, respectively (Fig. 2c and 2d, Supplementary Table S2). The results demonstrate the feasibility of the GNN-based deep learning model for cell-type classification at a single-cell resolution. scDeepSort were then systemically trained based on the HCL and MCA, on 562,977 human cells from 56 tissues and 201,746 mouse cells from 32 tissues.

### Performance comparison of scDeepSort with other methods

We collected human and mouse scRNA-seq data manually from high-quality studies, forming 76 external testing datasets (Supplementary Table S3) to comprehensively compare the performance of scDeepSort with other single-cell annotating methods. For the markers-dependent methods (CellAssign and Garnett), CellMatch was used as the reference for annotation. CellMatch is a comprehensive and tissue-specific cellular taxonomy reference database providing a panel of 20,792 human and mouse marker genes involved with 184 tissue types and 353 cell types. For profiles-dependent methods (SingleR and scMap), the HCL and MCA underlying scDeepSort were used as references. To ensure that all methods are able to predict the right cell identity, only cell types that were recorded in both cell marker database (CellMatch) and RNA-seq profiles (HCL and MCA) were selected to establish external testing datasets, which generated a total of 260,988 testing cells for human and mouse (Supplementary Table S4).

There are 27 human external testing datasets containing a total of 126,384 cells involving 10 tissue: blood, brain, colorectum, fetal kidney, kidney, liver, lung, pancreas, placenta and spleen. By mapping the predicted cell label with the real one (Supplementary Table S5), scDeepSort annotated testing cells with a 85.79% accuracy, higher than all other methods (CellAssign, 13.97%; Garnett, 26.27%; SingleR, 50.46%; scMap, 20.14%; Table 1). scDeepSort assigned the most accurate cell labels across 10 tissues and across 22 cell types (Fig. 4a and 4b) compared to other methods, the accuracies of which were mostly lower than 40%. Although the median accuracy of scDeepSort was 66.22%, its performance was significantly superior to other methods (Fig. 4c). At the level of cell type among 10 tissues, the results were strikingly concordant to the tissue-level results: the median accuracy of scDeepSort rose to 80.83%, whereas the median accuracies of CellAssign, SingleR and scMap were only 9.88%, 28.40% and 7.25% and that of Garnett was near 0% (Fig. 5d).

**Table 1.** Comparison of scDeepSort with other methods.

| Method |  | scDeepSort | CellAssign | Garnett | SingleR | scMap |
| --- | --- | --- | --- | --- | --- | --- |
|  |  | Markers-free | Markers-dependent | Markers-dependent | NA | NA |
|  |  | Profiles-free | NA | NA | Profiles-dependent | Profiles-dependent |
| Accuracy across external testing cells | Human (126,384) | 85.79% | 13.97% | 26.27% | 50.46% | 20.14% |
|  | Mouse (134,604) | 81.30% | 12.51% | 8.25% | 74.28% | 32.39% |
| Accuracy across tissues (median+SE) | Human | 66.22±7.98% | 11.25±5.18% | 9.82±4.50% | 18.61±6.52% | 9.26±4.14% |
|  | Mouse | 91.49±3.55% | 0.00±3.95% | 0.02±1.16% | 80.46±3.95% | 23.08±4.33% |
| Accuracy across cell types (median+SE) | Human | 80.83±4.48% | 9.88±3.36% | 0.43±2.63% | 28.40±3.86% | 7.25±3.66% |
|  | Mouse | 89.19±3.76% | 4.15±4.15% | 0.00±1.30% | 81.38±3.47% | 9.30±3.34% |

NA, not available.

**Fig. 4.**
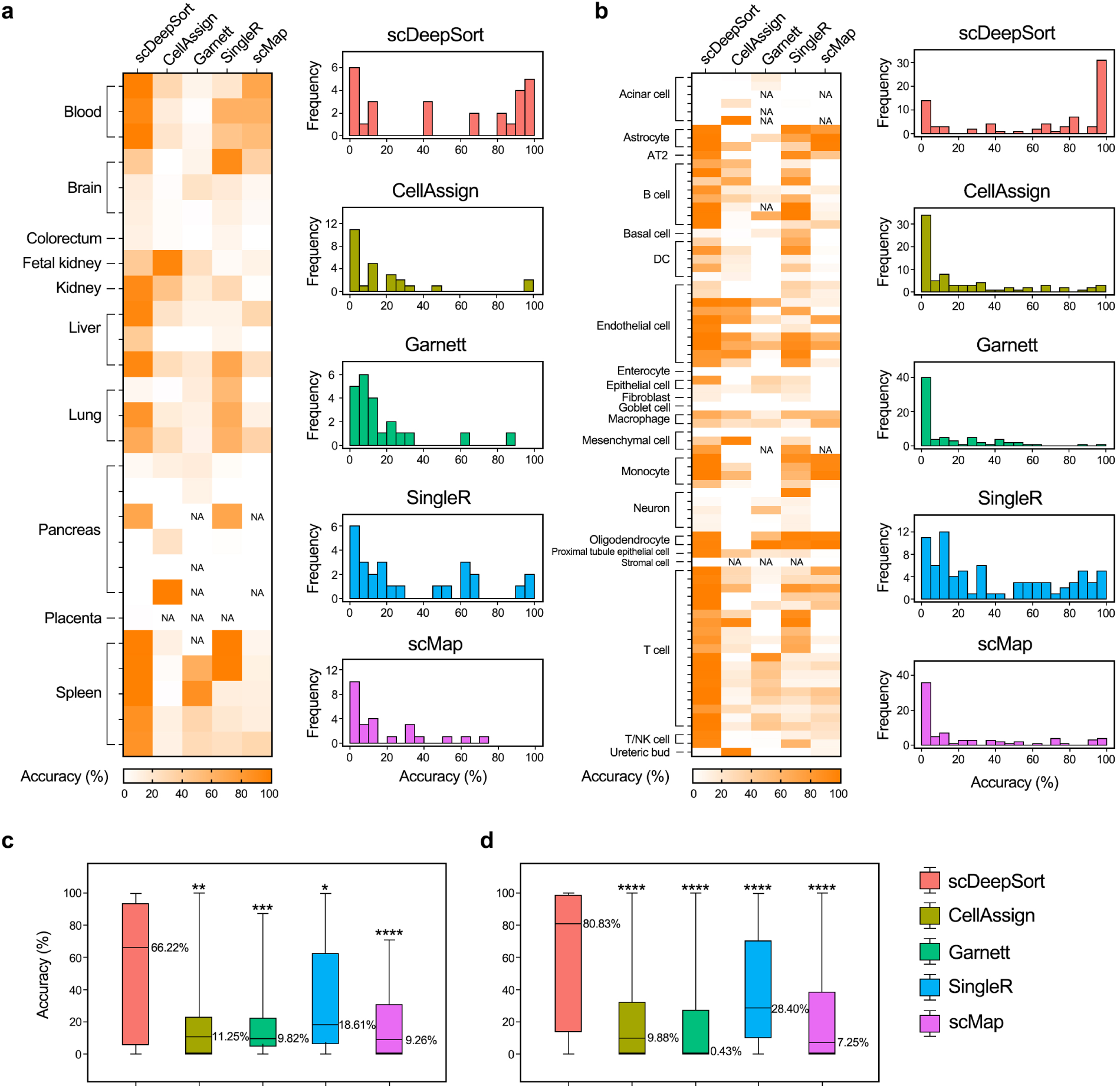
Performance comparison on human external testing datasets. **a)** Heatmap and distribution histogram of accuracies on 27 external testing datasets across 10 tissues by scDeepSort, CellAssign, Garnett, SingleR and scMap. NA, not available. **b)** Heatmap and distribution histogram of accuracies across 22 cell types. **c)** Boxplots summarized the maximal, minimal, median and quantile tissue-level accuracies for each method. The median accuracy is labelled beside the corresponding box. **d)** Boxplots summarized the statistical parameters at the level of cell types. Differences between multiple groups were determined using the matched ANOVA test by mixed-effects analysis with Dunnett’s multiple comparisons test (scDeepSort as control; *, *p* < 0.0332; **, *p* < 0.0021; ***, *p* < 0.0002; ****, *p* < 0.0001).

**Fig. 5.**
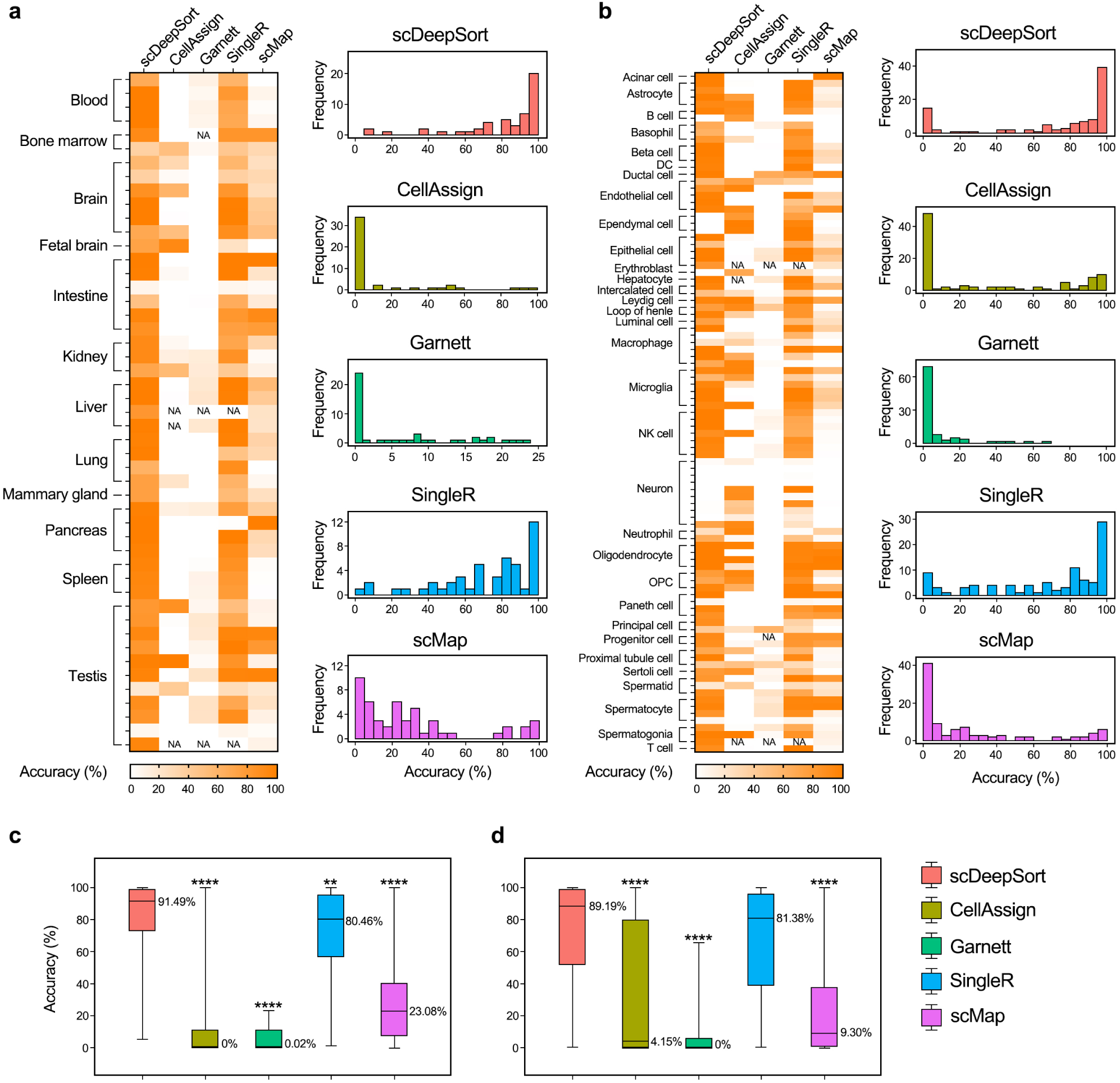
Performance comparison on mouse external testing datasets. **a)** Heatmap and distribution histogram of accuracies on 49 external testing datasets across 18 tissues by scDeepSort, CellAssign, Garnett, SingleR and scMap. NA, not available. **b)** Heatmap and distribution histogram of accuracies across 32 cell types. **c)** Boxplots summarized the maximal, minimal, median and quantile tissue-level accuracies for each method. The median accuracy is labelled beside the corresponding box. **d)** Boxplots summarized the statistical parameters at the level of cell types. Differences between multiple groups were determined using the matched ANOVA test by mixed-effects analysis with Dunnett’s multiple comparisons test (scDeepSort as control; **, *p* < 0.0021; ****, *p* < 0.0001).

Although scDeepSort outperformed other methods overall, it performed worse than other methods on some individual testing datasets at the level of tissues and cell types. For example, CellAssign obtained the highest accuracy among all methods on annotating the fetal kidney, the 6^th^ pancreas, the last acinar cell and the 2^nd^ mesenchymal cell datasets, etc. (Fig. 4a and 4b). On annotating the 1^st^ brain and the 3^rd^ pancreas datasets, SingleR perfectly classified the most cells’ identities (Fig. 5b). The accuracy of Garnett and scMap both reached 100% on the 2^nd^ oligodendrocyte dataset.

Surprisingly, all tested methods hardly predicted the accurate cell types on several human testing datasets, e.g., the 2^nd^ and the 3^rd^ brain, colorectum and most pancreas datasets (Fig. 4a), corresponding to the brain neuron, colorectum enterocyte and pancreas acinar datasets (Fig. 4b, Supplementary Table S5). For example, the 2,506 neurons in the 2^nd^ brain dataset^23^ were typically assigned as astrocytes by scDeepSort (92.98%) and scMap (73.34%), as neutrophils by CellAssign (39.55%), as unknown by Garnett (34.00%), and as T cells by SingleR (48.96%), whereas the 1,758 neurons in the 3^rd^ brain dataset^24^ were mainly classified as astrocytes by scDeepSort (91.81%) and scMap (75.14%), as macrophages by CellAssign (28.33%), as unknown by Garnett (67.12%), and as fetal enterocyte by SingleR (51.71%) (Supplementary Table S6). This result indirectly indicated the similarity between neurons and astrocytes, possibly because astrocytes can enter a neurogenic program by blocking Notch signaling^25^.

In the same manner, we also evaluated the performance of scDeepSort and other methods on annotating 49 external testing datasets of mouse cells, which includes 134,604 cells from 12 tissues: blood, bone marrow, brain, fetal brain, intestine, kidney, liver, lung, mammary gland, pancreas, spleen and testis. By mapping the predicted cell label with the real one (Supplementary Table S5), scDeepSort and SingleR accurately annotated the most cells for most of the external testing datasets (Fig. 5a and 5b). The median accuracies for scDeepSort and SingleR reached 91.49% and 80.46% at the tissue level (Fig. 5c). However, CellAssign and Garnett seemed unable to distinguish the most cells’ identities, accurately predicted only 12.51% and 8.25% cell identities across 134,604 mouse cells (Table 1). At the level of cell type among 12 tissues, the results were strikingly similar (Fig. 5b): the median accuracy of scDeepSort and SingleR still reached 89.19% and 81.38%, whereas the median accuracy of CellAssign and Garnett were both low, to about 0% (Fig. 5c and 5d). For scMap, it accurately predicted 32.39% of cell identities overall testing cells and its median accuracies were 23.08% and 9.30% at the levels of tissue and cell type, respectively (Table 1, Fig. 5c and 5d).

Interestingly, SingleR also performed well on the prediction task across the major testing datasets, realizing 74.28% accuracy over all testing cells, only a little worse than scDeepSort at 81.30% (Table 1). Although CellAssign and scMap performed poorly overall, they outperformed scDeepSort on some testing datasets. For instance, CellAssign obtained the highest accuracy among all methods in annotating the fetal brain dataset, the 1^st^ and 8^th^ testis datasets and 6^th^ macrophage dataset (Fig. 5a and 5b). In annotating the 2^nd^ B cell, erythroblasts, 2^nd^ microglia and 6^th^ neuron datasets, CellAssign perfectly classified the most cells’ identities (Fig. 5b). The accuracy of scMap achieved 100% in classifying the 28 cells in the 1^st^ intestine dataset and 108 cells in the 2^nd^ pancreas dataset, corresponding to paneth and acinar cells, respectively. Among mouse external testing datasets, Garnett’s accuracies were always low, ranging from 0% to 23.41% (Fig. 5a) and 0%-65.45% (Fig. 5b).

We note that all methods seldom predicted accurate cell types for some mouse testing datasets, e.g., the 3^rd^ intestine and 8^th^ and 11^th^ testis datasets. For example, the 3^rd^ intestine dataset comprised 260 intestine paneth cells marked by marker genes in the literature^26^. However, many of these cells were predicted to be epithelial cells by scDeepSort (92.69%), CellAssign (100%), SingleR (93.08%) and scMap (58.08%) (Supplementary Table S6), which may be unsurprising, as paneth cells are post-mitotic intestinal epithelial cells^27^.

In short, scDeepSort performed significantly better than all reference-dependent methods on 27 human datasets and 49 mouse datasets across 126,384 cells and 134,604 cells, respectively (Table 1). Some inconsistent predictions may have been caused by insufficient training samples, as in the case of human pancreas acinar cells (2 training cells), human liver mesenchyme cell (1 training cell), human pancreas mesenchyme cell (9 training cells), mouse bone marrow erythroblasts (11 training cells), mouse brain ependymal cells (9 training cells), and mouse pancreas B cells (16 training cells) (Supplementary Table S5). Besides, the transcriptomics similarity between transformable cells possibly lead to incorrect prediction, as shown in human brain neuron datasets. Moreover, unclear definition of cell types, subtypes and their relationship might cause imperfect mapping with the true cell identity, as the example of the intestine paneth cells described previously.

## Discussion

In this study, we developed a reference-free scalable cell-type annotation tool for single-cell transcriptomics data by using a deep learning model with a weighted GNN. From human and mouse scRNA-seq datasets, scDeepSort was able to be able to annotate most cells under the context of a specific organ. Moreover, scDeepSort significantly outperformed reference-based methods, i.e., the profiles-dependent CellAssign and Garnett and the markers-dependent SingleR and scMap. It is noted that the performance of our designed weighted GNN-based scDeepSort improves a lot in predicting cell types for most internal datasets compared to the traditional GNN-based deep learning model (Supplementary Table S2), indicating the superiority of our weighted GNN-based deep learning model in processing big data like high-throughput scRNA-seq data and in prediction. Moreover, the excellent performance of scDeepSort benefits from the recently published high-quality underlying data (HCL^8^ and MCA^21^) with the same scRNA-seq platform, which are the most comprehensive scRNA-seq data up to date across major tissues for human and mouse.

As reference-dependent methods, SingleR and scMap must compare cells with the reference of RNA-seq profiles, while CellAssign and Garnett need to be trained before annotating testing cells. Obviously, these reference-dependent methods are time-consuming, especially when using large reference databases. In addition, the increasing number of reference cell types and the corresponding markers or large RNA-seq reference profiles requires a high-quality processor and large memory capacity. However, scDeepSort realizes reference-free cell-type prediction, enabling fast annotation that does not require precise server configuration.

Because markers or RNA-seq profiles from different organs may vary considerably for the same cell type or be strikingly similar for different cell types^8,21,28^, another strength of scDeepSort is its comprehensive tissue-specific annotation, covering 56 human tissues and 32 mouse tissues. For example, the default references of SingleR are Encode^29^ and Blueprint Epigenomics^30^ for human cells and the Immunological Genome Project^31^ for mouse cells, without user-defined tissue types, which might increase incorrect cell identity predictions

Noted that there are some methods produced “not available” results when annotating external testing datasets. Specifically, CellAssign is not able to process some scRNA-seq data matrix containing cells with zero library sizes, and there may be too few training samples for some cell types at the root of a cell type hierarchy when training the classifier using Garnett. As for SingleR and scMap, an error may be present when executing hierarchical clustering on the SingleR scores and when fitting a linear model to select features for scMap. Apparently, these shortages tremendously limit the extension of these methods, which have not yet occurred when using scDeepSort.

Undoubtedly, scDeepSort’s performance depends on the underlying human and mouse single-cell transcriptomics atlases. Limited training datasets might influence cell-type annotation via scDeepSort, especially for these cell types without sufficient training data. However, future scRNA-seq studies will enable the expansion and perfection of atlases across the two species. Comprehensive integration of HCL, MCA and external testing datasets will greatly improve the performance of scDeepSort in turn.

Above all, the present results showed that scDeepSort can greatly help scientists sort single cells with an accurate cell label without prior reference knowledge, i.e., markers or RNA-seq profiles, significantly outperforming other popular annotation methods. scDeepSort realizes the reference-free tissue-specific cell-type annotation for single-cell transcriptomics data across two species by using comprehensive cell atlases, which may tremendously facilitate scRNA-seq studies and provide novel insights into mechanisms underlying biological process, as well as disease pathogenesis and progression.

## Acknowledgements

This work is supported by National Natural Science Foundation of China (81973701), the Natural Science Foundation of Zhejiang Province (LZ20H290002), and National Youth Top-notch Talent Support Program (W02070098).

## Author contributions

X.F. and H.C. conceived and designed the study. X.S., J.L., P.Y., Y.C. and X.L. collected and analysed the single-cell RNA-seq data. H.Y., X.Z. and Y.Y. implemented the algorithm of scDeepSort. X.S. and H.Y. developed the package of scDeepSort. All authors wrote the manuscript, read and approved the final manuscript.

## Methods

### Datasets

All scRNA-seq datasets were retrieved from several high-quality reports and the Gene Expression Omnibus (GEO), including human and mouse primary tissues, wherein unannotated cells were excluded and normal or healthy cells were included. The human cell landscape (HCL, https://figshare.com/articles/HCL_DGE_Data/7235471) provided data for 562,977 cells from 56 types of tissues and the mouse cell atlas (MCA, https://figshare.com/articles/MCA_DGE_Data/5435866) provided 201,764 cells involving 32 tissues. External testing datasets used for comparing scDeepSort with other methods were freely available from public platforms detailed in Supplementary Table S3.

### Data preprocessing

All scRNA-seq data were preprocessed using R (version 3.6.1). For the Zheng dataset, the raw count was processed in accordance with the pipeline detailed in the Satija Lab tutorial, using Seurat 3.0, wherein cells with more than 2,500 or fewer than 200 unique features or with mitochondrial counts greater than 5% were filtered out. For other datasets, all cells in the datasets were included in the filtered matrices. Human and mouse gene symbols were revised in accordance with NCBI gene data (https://www.ncbi.nlm.nih.gov/gene/) updated on Jan. 10, 2020, wherein unmatched genes and duplicated genes were removed. For all human and mouse datasets, the raw data were normalized via the global-scaling normalization method LogNormalize in preparation for running the subsequent scDeepSort pipeline and other methods.

### scDeepSort algorithm

scDeepSort consists of three components: the embedding layer, weighted graph aggregator and linear classifier layers. The embedding layer stores the representation of graph nodes and is freezed during training. The weighted graph aggregator layer inductively learns graph structure information, generating linear separable feature space for cells. In this layer, a modified version of the GraphSAGE information processing framework was applied as the backbone GNN. The final linear classifier layer classifies the final cell state representation produced from the weighted graph aggregator layer into one of the predefined cell type categories.

#### Weighted cell-gene graph generation

To construct the weighted cell-gene graph, cells and genes were both treated as graph nodes and the gene expression for each cell was regarded as the weighted edge between cells and genes, constituting the embedding layer. First, we used dimensionality reduction methods to obtain the node embeddings for cells and genes. For an input single-cell data matrix *D* ∈ ℝ^*m*×*n*^ (*m* genes and *n* cells), principal component analysis (PCA) was applied to extract dense representations of a fixed-size dimension (*d* = 400) as gene representations. A weighted sum of gene representations with single-cell data matrix *D* as input was used to obtain the cell representations with the same dimension *d*. By collecting gene and cell representations, a matrix *X* ∈ ℝ^(*m*+*n*)×*d*^ was constructed as the initial node embeddings. Second, a weighted adjacency matrix 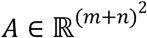 was generated from the input single-cell data matrix *D*, in which the gene expression (> 0) was directly regarded as the weights of edges between cells and genes.

#### Aggregating process

To inductively learn graph structure information, we followed a graph neural network framework called GraphSAGE. The essential processes of GraphSAGE are sampling a batch of 500 nodes with their neighbors and aggregating graph neighborhood to generate node representations for each node. However, we proposed a new weighted graph aggregator layer to replace the aggregator of GraphSAGE. Let 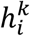 (a 200-dimensional vector in our experiments) represents the embedding of node *i* in the *k*^th^ layer. Our weighted graph aggregator layer can be summarized as:

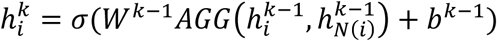

where *N*(*i*) is the set of one-hop neighbors of node *i*. The output of the aggregate function AGG is then transformed to target dimension by a linear transformation shared among all nodes, followed by a non-linear activation function σ called Rectified Linear Unit (ReLU). In practice, we set *k* = 1. The aggregate function *AGG* contains two newly techniques. The first technique is called weighted adjacency matrix normalization. The main reasons for applying normalization to weighted adjacency matrix are twofold. Gene expression varies a lot across different kinds of cells. For single cell, the expression level and pattern of different genes can also vary. Thus, we normalize weighted adjacency matrix *A* as following:

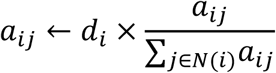

where *a*_*ij*_, the weight of an edge from node *j* to node *i*, is the element of *A*, and *d*_*i*_ denotes the indegree of cell node *i*. The second technique is the learnable sharing confidence. Due to batch effect and missing value issues, we proposed to add learnable parameters to each edges as a confidence matrix while leveraging the context of one-hop neighborhood of nodes in a weighted graph. For a gene node *j*, we proposed a learnable sharing parameter *β*_*j*_ as the confidence value for the edges that interact with node *j*. Another learnable parameter *α* as the confidence value of the self-loop edge for each cell. Its value will be shared among cells since we may encounter with new cells in testing time. Therefore, the overall formulation of gathering neighborhood information given each sub-graph of cell node *i* is stated below:

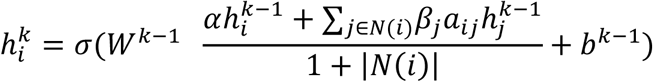

#### A linear classifier layer

The weighted graph aggregator layer produces a latent feature space for the graph. To classify the final cell state representation into one of the pre-defined cell-type categories, we extract cell node representations and feed them into a linear classifier layer.

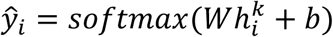

Cross entropy loss was then used to measure the difference between the predicted class distribution and the labels. Therefore, the objective function can be written as:

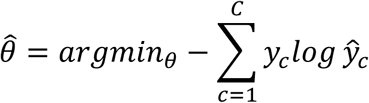

We train our model with the above objective function using a stochastic gradient descent method called Adam, with default hyper parameters except for the learning rate of 0.001 and the weight decay rate of 0.0005 until convergence or after 500 epochs.

### scDeepSort performance evaluation on internal datasets

For each cell type, cells numbering at least more than 5‰ of the total cells in each tissue were included and randomly divided into training and testing sets, ensuring that the ratio of training and testing cells was set to 8:2. For each tissue from the human and mouse atlases, all training cells of various types were merged and supervised learned with the GNN-based deep learning model for cell-type prediction on the testing cells originated from the same tissue.

### Performance comparison with other methods on external testing datasets

CellMatch, MCA or HCL were used as the reference datasets for reference-dependent methods. In order to compare the performance of scDeepSort with other methods on annotating cell types of single-cell transcriptomics data, only the cell types that existed in both cell marker database (CellMatch) and RNA-seq profiles (MCA and HCL) were selected to construct the testing datasets.

For CellAssign, external testing datasets were first transformed as SingleCellExperiment objects with a normalized matrix. The CellMatch database containing tissue-specific cell markers was then used as reference. All other parameters in CellAssign were kept as default (i.e., the learning rate was set to 0.01).

For Garnett, marker genes from the CellMatch database were extracted and checked to train classifiers for each testing dataset. The parameter of the number of unknown type cells was set as 50 during classification. Then, the trained classifiers were used to classify the cells for each test dataset.

For SingleR and scMap, external testing datasets were transformed into SingleR and SingleCellExperiment objects and annotated based on reference database of scRNA-seq profiles. To annotate human and mouse testing datasets, scRNA-seq profiles from human and mouse cell atlases were used as the reference database for human and mouse, respectively.

### Accuracy evaluation

For scDeepSort, CellAssign, SingleR and scMap, accuracy is defined as the percentage of consistent cells with the same cell type, as in the literature.

### Data availability

No new data was generated for this study. All data used in this study is publicly available as previously described.

### Code availability

scDeepSort is available as a python package (https://github.com/ZJUFanLab/scDeepSort) and the source code and results of comparison with other methods are available at github (https://github.com/ZJUFanLab/scDeepSort_performace_comparison).

### Statistics

R (version 3.6.1) and GraphPad Prism 8.0.1 were used for the statistical analysis. Differences between multiple groups were determined using the matched ANOVA test by mixed-effects analysis with Dunnett’s multiple comparisons test (significant with *p* < 0.0332).

